# *Monanthotaxis bali* (Annonaceae) a new Critically Endangered (possibly extinct) montane forest treelet from Bali Ngemba, Cameroon

**DOI:** 10.1101/2022.07.04.498636

**Authors:** Martin Cheek, Iain Darbyshire, Jean Michel Onana

## Abstract

*Monanthotaxis bali* is the only known, solely montane (occurring solely above 2000 m alt.) species of the genus. It joins *Monanthotaxis orophila* (Rwanda) and *M. discolor* (Tanzania), two other species that can also occur above 2000 m alt. *Monanthotaxis bali* is an addition to the small number (28) of the tree species of the surviving montane forests of the Cameroon Highlands of which only eight other species are endemic.

Due to its supra-axillary inflorescences, and petals arranged in one whorl but with the outer petals overlapping the inner petals distally, and rounded flower buds, the new species is placed in Hoekstra’s clade B of *Monanthotaxis*. The new species is unusual in being a treelet in a predominantly lianescent genus, and in lacking the glaucous underside of the leaf-blades that usually characterises the genus. *Monanthotaxis bali* takes the number of continental African species of the genus to 80, and makes Cameroon, with 30 species, the most species-diverse country for the genus.

*Monanthotaxis bali* is known only from the Bali Ngemba Forest Reserve, a forest remnant under pressure of degradation and clearance in the Bamenda Highlands of Cameroon. It may already be extinct due to logging and agricultural activities. Here it is described, illustrated, mapped and assessed as Critically Endangered (Possibly Extinct) using the IUCN 2012 criteria.

## Introduction

In Jan. 2022, whilst examining herbarium specimens collected for the Conservation Checklist of the Plants of Bali-Ngemba, Cameroon (Harvey *et al*. 2004) but which had by oversight not been identified, an unknown taxon of Annonaceae was encountered. *Darbyshire* 375 was collected 11 April 2004 (YA, K) at 2110 m alt. This is an extremely high altitude for a member of Annonaceae to occur in the Cameroon Highlands. On the higher mountains, Mt Cameroon (4095 m alt) and Mt Oku (3011 m alt.), the highest recorded occurrences of Annonaceae are respectively at 1400 m (*Uvariodendron fuscum* (Benth.) R.E. Fries and 1500 m alt. *Monanthotaxis vulcanica* P.H. Hoekstra (as *M. littoralis* (Bagsh. & Bak.f.) Verdc. (Cable & Cheek 1998; Cheek *et al*. 2000).

*Darbyshire* 375 was identified as a *Monanthotaxis* Baill. in the pantropical tribe Uvarieae, despite being a treelet with leaf-blades yellow-brown on the abaxial surface, when this genus is recognised by its lianescent habit and glaucous lower side of the leaves (Hoekstra *et al*. 2021). Pointers to placement in *Monanthotaxis* are the presence of a bract midway along the pedicel, the supra-axillary inflorescences and above all, the presence of dark coloured, gland-like structures at the base of the leaf blade, a feature almost exclusively present in *Monanthotaxis* and not found in other Annonaceae (Hoekstra *et al*. 2021). The Annonaceae is a pantropical family largely restricted to tropical rainforest habitats. The characteristic often showy flowers and fruit help make it widely collected and studied (Chatrou *et al*. 2012; Couvreur *et al*. (in press)). Annonaceae are well known for species producing edible fruits (bullocks heart, soursop, custard apple), and for the fragrant ylang-ylang (*Cananga odorata* (Lam.) Hook.f. & Thomson) used in perfumes. The Annonaceae is a basal angiosperm family in the order Magnoliales (Heywood *et al*. 2007). It comprises c. 2400 species and c. 107 genera of trees, shrubs and lianas (Guo *et al*. 2017a). Within the subfamily Annonoideae comprising over 1500 species, the pantropical tribe Uvarieae is composed of 15 genera and 474 species (Guo *et al*. 2017a).

New Annonaceae taxa are continually being discovered in Africa, including new genera to science e.g. *Lukea* Cheek & Gosline in E Africa (Cheek *et al*. in press) and new species, e.g. of *Uvariodendron* (Dagallier *et al*. 2021) also in E Africa, and in Cameroon, new species such as *Uvariopsis dicaprio* Cheek & Gosline (Gosline *et al*. 2022), *U. etugeana* Dagallier & Couvreur (Couvreur et al. in press) and *Xylopia monticola* D.M. Johnson & N. A. Murray (Johnson & Murray 2018)

The genus *Monanthotaxis* is confined to subsaharan continental Africa (79 species), the Comores and Madagascar (Hoekstra *et al*. 2021). The numbers of species have been recently increased by the description of about 20 new species to science (Hoekstra *et al*. 2014; 2016; 2021) and the subsumation of the African species of the genera *Friesodielsia* Steenis, *Exellia* Boutique and *Gilbertiella* Boutique (Guo *et al*. 2017b). Earlier, Verdcourt (1971) had subsumed the African species of *Popowia* Endl. and *Enneastemon* Exell into *Monanthotaxis*, recognising 55 species in the last genus.

*Monanthotaxis* species are mainly confined to rainforest, with only a few species occurring in woodland, notably the widespread *M. buchananii* (Engl.) Verdc. In continental Africa species extend from Senegal in the north, and west to Somalia in the east, and as far south as the Eastern Cape Province of South Africa (Hoekstra *et al*. 2021).

Fifty four species occur in the Central African forest, with DR Congo and Cameroon both having the highest number of all countries: 29 species. Only a few forest species have wide distributions e.g. *Monanthotaxis laurentii* (De Wild.) Verdc. from Guinea in the west to eastern DR Congo in the east (Hoekstra *et al*. 2021). Hoekstra *et al*. (2021) identify nine different clades, A-I, based on 5 plastid and 2 nuclear DNA markers. Each clade is morphologically characterised. Seed dispersal is mainly recorded to be by primates including humans, gorillas, chimpanzees, mandrills and galagos in continental Africa, and in Madagascar, lemurs. The fruits are juicy, sweet and often brightly coloured red or orange (Hoekstra *et al*. 2021).

Several species are used in traditional medicine including as anti-malarials, and 15 species have been studied for their phytochemistry (Hoekstra *et al*. 2021).

## Materials & methods

The specimen was collected using the patrol method as documented in Cheek & Cable (1997). Unambiguous identification as a new species was largely thanks to the excellent monograph including comprehensive species delimitations and keys, of Hoekstra *et al*. (2021) without which this paper would have been much more difficult to complete so speedily. Herbarium material was examined with a Leica Wild M8 dissecting binocular microscope fitted with an eyepiece graticule measuring in units of 0.025 mm at maximum magnification. The drawing was made with the same equipment with a Leica 308700 camera lucida attachment. Specimens, or their high-resolution images, were inspected from the following herbaria: BR, K, P and WAG. All specimens cited have been seen. Names of species and authors follow the International Plant Names Index (IPNI continuously updated). Nomenclature follows Turland *et al*. (2018). Technical terms follow Beentje & Cheek (2003) and Hoekstra *et al*. (2021). The conservation assessment follows the IUCN (2012) categories and criteria. Herbarium codes follow Index Herbariorum (Thiers continuously updated).

## Results

Because *Darbyshire* 375 has bisexual flowers, supra-axillary 1 – 2-flowered inflorescences, 7 – 9 carpels, 9 free stamens, no staminodes, inner petals visible between the outer petals in the rounded flower buds, hairy petals, ovules 4 – 6, hairs on young branches yellow-brown, it keys out in Hoekstra *et al*. (2021) to couplet 45 which leads to *Monanthotaxis barteri* (Baill.) Verdc. of West Africa and *M. schweinfurthii* (Eng. & Diels) Verdc. (Central Africa). It differs from these two species in numerous characters (see Table 1 and diagnosis below).

**Table 1.** Characters separating *Monanthotaxis barteri* and *M. schweinfurthii* from *M. bali*. Data for the first two species from Hoekstra *et al*. (2021) and herbarium specimens at K.

|  | *Monanthotaxis barteri* | *Monanthotaxis bali* | *Monanthotaxis schweinfurthii* |
| --- | --- | --- | --- |
| Habit | Liana, lianescent shrub, rarely ‘a shrub with spreading branches’ | Treelet | Liana, lianescent shrub, rarely a shrub |
| Leaf surfaces & texture | Discolorous, lower surface glaucous white or white-grey, upper surface green or grey-green. Chartaceous or thinly coriaceous | Concolorous, both surfaces pale brown. Coriaceous | Discolorous, lower surface glaucous, upper surface green. Chartaceous or thinly coriaceous |
| Petiole length (mm) | 2.5 – 6 | (5 – )6 – 9( – 12) | 4 – 9 |
| Flower bud diam. (mm) | 2 – 3.5 | (5 – )7 – 9 | 3 – 4.5 |
| Pedicel diam. (mm) | 0.25 – 0.3 | 0.5 – 0.75( – 1.1) | 0.3 – 0.4 |
| Inflorescence insertion | Axillary (arising next to the leaf petiole) | Supra-axillary (arising 1 mm above the leaf axil). | Axillary |
| Stamen length (mm) & indumentum | 1.2 – 1.4, hairy | c. 2, glabrous | 1 – 1.7, hairy |
| Anther thecae position & shape of stamen apex | Extrorse, co-adjacent; stamen apex flat. | Latrorse, thecae separated, stamen apex with depression (between the thecae) | Extrorse, co-adjacent; stamen apex flat projecting over the anthers |
| Outer petals dimensions (mm) | 3.2 – 3.6( – 5) x 2.2 – 2.6( – 3.4) | 7.3 – 7.6 x 7.1 – 7.3 | 4 – 4.8 x 2.8 – 4.6 |
| Inner petal dimensions (mm) | 2.7 – 3.1( – 4.5) x 1.5 – 2.3 | 6.7 – 7 x 3.6 – 4 | 2 – 4.3 x 1 – 3.3 |
| Altitudinal range (m) | 0 – 1000 | c. 2100 | 200 – 1290 |

*Monanthotaxis barteri* and *M. schweinfurthii* are placed in Hoekstra’s clade B together with *M. seretii* (De Wild.) P.H. Hoekstra, *M. aestuaria* P.H. Hoekstra, *M. ochroleuca* (Diels) P.H. Hoekstra, *M. foliosa* (Engl. & Diels) P.H. Hoekstra, *M. capea* (B.G. Camus & A. Camus) Verdc., and *M. biglandulosa* (Boutique) P.H. Hoekstra. This group of species is characterised by rounded flower buds in which the inner petals are exposed between the bases of the outer petals. They also have a single whorl of both stamens (usually 9) and carpels (usually 6 – 9), are supra-axillary (the inflorescences not arising immediately from the axil but some distance above it) and 1 – 2-flowered. This group, Clade B, equates to the formerly accepted genus *Enneastemon* Exell.

Here we formally describe *Darbyshire* 375 as new to science:

### **Monanthotaxis bali** *Cheek* **sp. nov**. Type

Cameroon, Northwest Region, Mezam Division, Bali Ngemba Forest Reserve, ridge at top of reserve, from Ntanyam Trail, 2110 m alt., fl. 11 April 2004, *Darbyshire* 375 with Nukam, Ronsted, Atem (holotype K, barcode K000593350, isotypes US, YA). (Fig.1 &2)

**Fig. 1.**
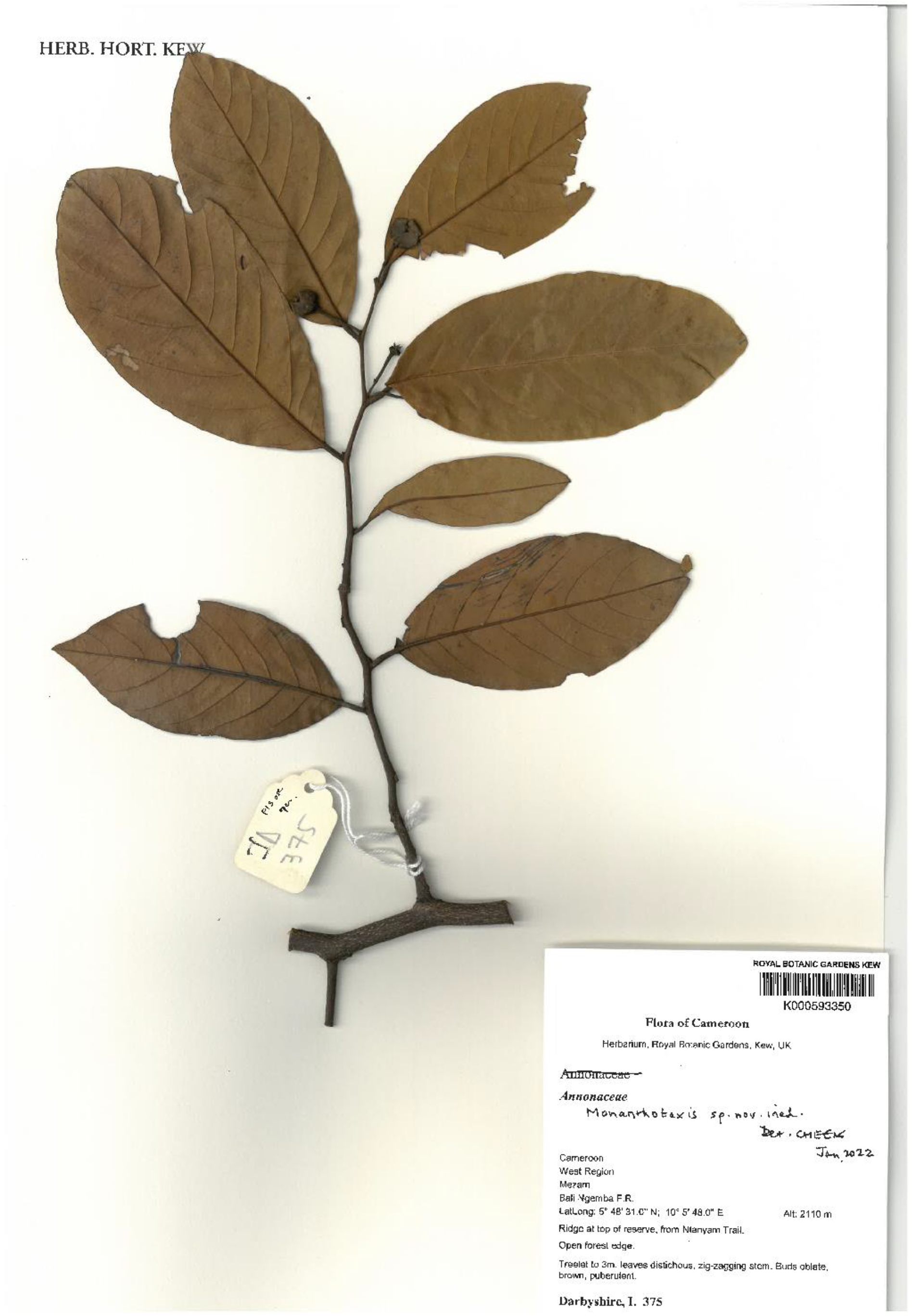
*Monanthotaxis bali* The type specimen *Darbyshire* 375 (holotype K, barcode K000593350).

**Fig. 2.**
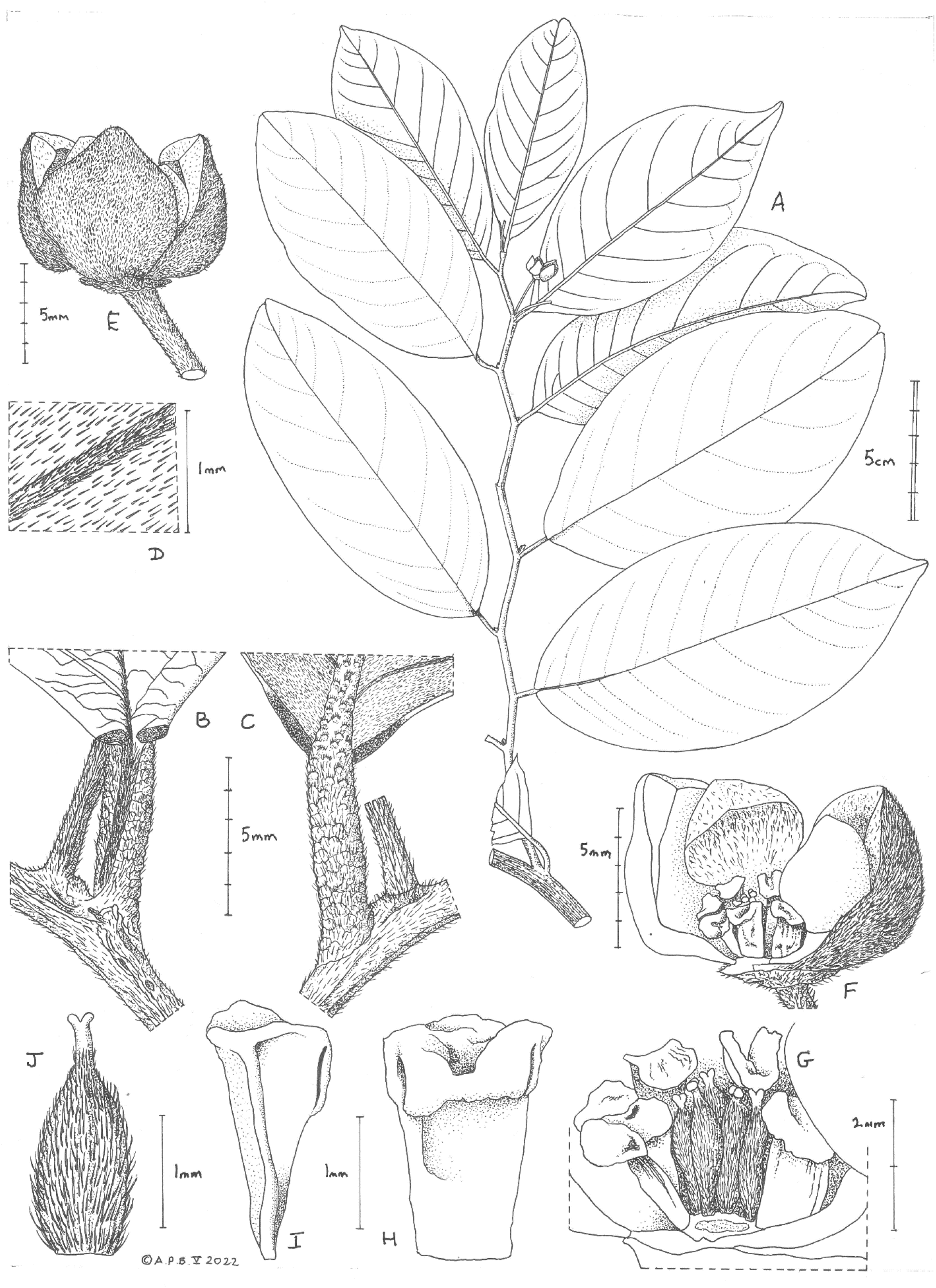
*Monanthotaxis bali* **A** habit, flowering stem; **B** node showing supra-axillary inflorescence base and paired glandular area at base of adaxial leaf blade; **C** as B but abaxial surface of blade; **D** indumentum on abaxial leaf blade surface, including secondary vein; **E** flower, side view; **F** flower, after removal of an inner and outer petal to show androecium and gynoecium; **G** close-up of F, showing detail of carpels after removal of stamen; **H** stamen outer face, showing the two flanking thecal aperture and central apical depression; **I** stamen as H, but side view, showing absence of hairs; **J** pistil. All drawn from *Darbyshire* 375 by ANDREW BROWN.

*Treelet* c. 3m tall, branches with leafy stems flexuose (zig-zag), terete, 6 – 7 mm diam., dull black with numerous low longitudinal ridges, lenticels conspicuous bright white orbicular (-elliptic) 0.5 – 0.75 × 0.25 – 0.5mm, glabrous. *Leafy stems* distichous, flexuous, terete, 1.5 – 2.5 mm diam., internodes 1.5 – 2.5(– 3) cm long, dark purple-brown, first internode 70 – 80% covered in appressed translucent or slightly yellow simple hairs 0.2 – 0.25 mm long, eventually glabrescent, at second node surface about 50% covered. *Leaves* leathery, drying glossy pale yellow brown above, matt yellow-brown below, not glaucous, elliptic, elliptic-oblong or elliptic-obovate (6 –)9.3 – 13.7(– 15) x (4.0 –)4.4 – 6.8(– 7.3) cm, shortly acuminate or rounded at apex, base obtuse-rounded or rounded and minutely cordate, the lobe margins glossy-black, gland-like, 0.3 – 0.75 × 1mm, midrib and secondary nerves impressed on adaxial surface, raised on the abaxial surface, eucamptodromous, secondary nerves 7 – 11(– 12) on each side of the midrib, arising at 45 – 60° from the midrib, gradually arching upwards, becoming parallel to the margin, percurrent, tertiary nerves inconspicuous abaxially, but with quaternary nerves highly prominent, forming a fine reticulum adaxial surface, with cells 0.25 – 0.5 mm diam., margin slightly revolute, adaxial surface densely white hairy in young leaves, soon glabrescent, then punctate, only the base of the midrib grey-hairy, abaxial surface with persistent appressed translucent or red hairs, scattered evenly covering 20 – 30% of the surface, nerves more sparsely hairy. Petioles grey-black, twisted through 90°, distal part canaliculate, gradually becoming terete proximally, (5 –)6 – 9(– 12) x 1.5 mm, sparsely appressed hairy, hairs as stem, articulation sub-basal, leaving an orbicular, cupular scar raised c. 0.5mm above the stem.

*Inflorescences* supra-axillary, arising c. 1 mm above the leaf axil, 1 or 2-flowered, pedicels drying black, shortly connate, proximally terete, (6 –)8 – 14(– 15) x 0.5 – 0.75(– 1.1) mm, lower bract absent, upper bract inserted slightly below the midpoint, clasping the pedicel for c. ⅓ the circumference, conical (by involution of margins) 0.5 – 1.25 mm long, inclined forward at c. 45° from pedicel axis, densely covered in translucent hairs as stem. Flower buds oblate in bud, brown (0.5 –)7 – 9 mm diam. *Flowers* bisexual, sepals connate, forming a shallowly cupular calyx 0.8mm x 4 mm, lobes not divided, apex rounded, indumentum densely pubescent on outer surface, hairs erect, crisped, translucent, c. 0.1 mm long, inner surface glabrous, drying purple, receptacle c. 3 mm diam., flat; petals brown at anthesis (collection notes), 6, base of inner petals visible in bud, outer petals overlapping at apex. *Outer petals* 3 subrhombic, thick, 7.3 – 7.6 × 7.1 – 7.3 mm, apex obtuse, base slightly united with inner petals, outer and inner surface densely hairy as the sepals, excepting a small glabrous area 2 – 2.5 mm diam. on adaxial surface at base. *Inner petals* 3 cucullate (hooded) in the upper half c. 4 × 4 mm, lower half broadly stipitate c. 3 × 2.5 mm, indumentum as outer petals. Stamens 9, in one whorl, free, 2 × 0.75 – 1.25 mm, filaments centripetally flattened, 1.25 – 1.5 × 0.75 – 1.25 mm, drying yellow, thecae latrorse, separated by a distal depression, connective prolonged horizontally inwards towards the carpels, flat, oblong, apex rounded c. 1.25 × 0.75 mm, glabrous, staminodes absent. Carpels 7 – 9 in one whorl, erect, narrowly ellipsoid c. 2 × 0.75 mm, densely covered in appressed 0.2 – 0.25 mm long simple white or yellow hairs, ovules 3-4, style-stigma erect cylindric c. 0.5 × 0.08 mm, slightly angled, stigma capitate, glabrous. Fruits not seen.

### RECOGNITION

Differing from *Monanthotaxis barteri* (Baill.) Verdc. and *M. schweinfurthii* (Engl. & Diels) Verdc. in being a treelet with concolorous leaves, supra-axillary inflorescences and flower buds 7 – 9 mm diam. (vs lianas with blades glaucous below, axillary inflorescences and flower buds 2 – 4.5 mm diam.). Additional diagnostic characters are given in Table 1.

### DISTRIBUTION

Cameroon, North West Region (formerly Province), Mezam Division, Bali Ngemba Forest Reserve, only known from remnants of montane forest.

### SPECIMENS EXAMINED: CAMEROON

Northwest Region, Mezam Division, Bali Ngemba Forest Reserve, ridge at top of reserve, from Ntanyam Trail, 2110 m alt., fl. 11 April 2004, *Darbyshire* 375 with Nukam, Ronsted, Atem (holotype K, barcode K000593346; isotypes US, YA).

### HABITAT

Infrequent treelet of montane evergreen forest remnant on basaltic plateau; c. 2110 m alt.

### CONSERVATION STATUS

Although within the Bali Ngemba Forest Reserve, *Monanthotaxis bali* is not protected because resources for patrolling seem so inadequate that people enter the forest to cut timber and plant food crops with impunity. Much of the montane, upper part of the reserve has been entirely cleared of forest for cultivation (Harvey *et al*. 2004), so it is fortunate that this single plant survived in one of the scraps that remained in 2004. Given continual pressure in the last 18 years it may be that this fragment has also been lost.

This distinctive treelet has not been found in surveys elsewhere in the Cameroon HIghlands and adjacent areas (Cheek 1992; Cheek *et al*. 1996; Cable & Cheek 1998; Cheek *et al*. 2000; Maisels *et al*. 2000; Chapman & Chapman 2001; Cheek *et al*. 2004; Harvey *et al*. 2004; Cheek *et al*. 2006; Cheek *et al*. 2010; Harvey *et al*. 2010; Cheek *et al*. 2011). Therefore, it may be, or have been, endemic to Bali Ngemba. Several other plants of high altitudes in the Cameroon Highlands appear to be point endemics e.g. *Impatiens etindensis* Cheek & Eb. Fisch. (Balsaminaceae, Cheek & Fischer 1999), *Ledermanniella pollardiana* Cheek (Podostemaceae, Cheek & Ameka 2008) and *Kupeantha fosimondi* (Cheek) Cheek (Rubiaceae), Cheek *et al*. 2018a). However, others, originally thought to have been point endemics have since been found at one or more other locations in the highlands of Cameroon, e.g. *Coffea montekupensis* Stoffelen (Rubiaceae, Stoffelen *et al*. 1997) and *Impatiens frithii* Cheek (Balsaminaceae, Cheek & Csiba 2002). It is to be hoped that the second scenario will be the case for *Monanthotaxis bali* and that it will also be found at other locations, although we consider this unlikely.

Bali Ngemba is less than 8 km^2^ in area, and it was relatively well-surveyed by large teams of botanical collectors 2000 – 2004 (Harvey *et al*. 2004). Other new, or rediscovered species from Bali-Ngemba are known from multiple gatherings, sometimes 10 or more e.g. *Leptonychia kamerunensis* Engl. & K. Krause (Byttneriaceae, Cheek *et al*. 2013), *Vepris onanae* Cheek (Cheek *et al*. 2022a). That only a single collection is known for *Monanthotaxis bali*, and that a single plant was reported (*Darbyshire* 375) suggests that the species is genuinely infrequent, if indeed it survives.

Here we assess *Monathotaxis bali* as Critically Endangered (Possible Extinct), CR (PE) B1+B2ab(iii) +D since only a single individual was reported from a forest scrap, threats are ongoing, and AOO and EOO are estimated as 4 km^2^ using the cells of that size preferred by IUCN. It has been estimated that 96.5% of the former area of the montane and submontane forests of the Bamenda Highlands has been lost to clearance (Cheek *et al*. 2000). The surviving forests of the Bamenda Highlands are extremely fragmented. The indications for seed dispersal in *Monanthotaxis* are that it is conducted by fruit-eating primates (e.g. chimpanzees) (Hoekstra *et al*. 2021) for which traversal of human-created habitat (farmland and secondary grassland) in the Cameroon Highlands is known to be problematic (Cheek *et al*. 2021a).

### ETYMOLOGY

Named for the Bali Ngemba Forest Reserve near the town of Bali.

### NOTES

Additional plant taxa unique to Bali Ngemba forest and sometimes also the nearly adjoining and smaller Baba 2 community forest are: *Vepris bali* Cheek (Rutaceae, Cheek *et al*. 2018b), *Psychotria babatwoensis* Cheek (Rubiaceae, Cheek *et al*. 2009), *Magnistipula butayei* subsp. *balingembaensis* [Sothers](https://www.ipni.org/a/39295-1), [Prance](https://www.ipni.org/a/12747-1) & [B.J.Pollard](https://www.ipni.org/a/39322-1) (Chrysobalanaceae, Pollard *et al*. in Harvey *et al*. 2004). Newly described endangered, tree species from Bali Ngemba include *Tricalysia elmar* Cheek (Rubiaceae, Cheek *et al*. 2020a), *Vepris onanae* Cheek (Cheek *et al*. 2022a) and *Deinbollia onanae* Cheek (Cheek *et al*. 2021a).

Sadly, *Monanthotaxis bali* is not the only seemingly point endemic in the genus in Africa. Eight other species are known from a single collection or location: *Monanthotaxis aquila* P.H. Hoekstra (Ivory Coast), *M. atewensis*. P.H. Hoekstra (Ghana), *M. couvreurii* P.H. Hoekstra (Cameroon), *M. filipes* P.H. Hoekstra (Tanzania), *M. glomerulata* (Le Thomas) Verdc. (Gabon, last seen 80 years ago), *M. hexamera* P.H. Hoekstra (Zingui, Cameroon, last seen 50+ years ago), *M. mortehanii* (De Wild.) Verdc. (DR Congo, last seen >100 years ago) and *M. zenkeri* P.H. Hoekstra (Bipinde, Cameroon, last seen 100 years ago). All of these species, completely logically, are treated as Critically Endangered by Hoekstra *et al*. (2021). *Monanthotaxis bali* of Cameroon occurs within the Cross-Sanaga Interval (Cheek *et al*. 2001) which has the highest species and generic diversity per degree square in tropical Africa (Barthlott *et al*. 1996; Dagallier *et al*. 2020) including endemic genera such as *Medusandra* Brenan (Peridiscaceae, Breteler *et al*. 2015; Soltis *et al*. 2007). Much of this diversity is associated with the Cameroon Highland areas, different highlands each having a species of a genus e.g. as in *Kupeantha* Cheek (Rubiaceae, Cheek *et al*. 2018a). However, unlike for many genera in Africa, such as *Vepris* (Cheek *et al*. 2022a) and *Impatiens* (Cheek *et al*. 2022b), the Cross-Sanaga is not notable as a centre of diversity for the genus *Monanthotaxis*, since it holds only 14 species of the genus, of which three are endemic to the interval, increasing to four with the publication of *Monanthotaxis bali*.

In tropical Africa Annonaceae predominantly occur in lowland evergreen forest. A remarkable feature of *Monanthotaxis* is that 33 of the 79 species documented by Hoekstra *et al*. (2021) can occur over 1000 m alt., while four other species occur only above 1000 m alt, two others occurring sometimes above 2000 m (see introduction).

## Discussion and Conclusions

*Monanthotaxis bali* joins the growing list of African species considered likely to be extinct. Species considered extinct in Cameroon are *Oxygyne triandra* Schltr., *Afrothismia pachyantha* Schltr. (Cheek & Williams 1999, Cheek *et al*. 2018c, Cheek *et al*. 2019). It seems possible that *Monanthotaxis bali* became extinct before it is formerly named scientifically in this paper, as was the case with *Vepris bali*, also of higher altitudes in Bali Ngemba (Cheek *et al*. 2018b). This extinction crisis extends through W. Africa e.g. *Saxicolella deniseae* Cheek, also seemingly extinct before publication (Cheek *et al*. 2022c). Similarly, *Pseudohydrosme bogneri* Cheek & Moxon-Holt and *P. buettneri* Engl. are now considered extinct in neighbouring Gabon (Moxon-Holt & Cheek 2020; Cheek *et al*. 2021b).

About 2000 species of vascular plant have been described as new to science each year for the last fifteen years or more. Cameroon currently has the highest number of new species to science published each year (Cheek *et al*. 2020b), recent examples being published are (Achoundong *et al*. 2021; Aguirre-Alvarez *et al*. 2021; Cheek *et al*. 2021a; 2021b; 2021c; Cheek & Onana 2021; Couvreur *et al*. in press; Gosline *et al*. 2021). While some exceptions exist (Cheek & Etuge 2009; Cheek *et al*. 2019), many, probably most of these species are highly range-restricted, making them especially at risk of habitat destruction, and so likely to be rated as threatened when assessed.

Only 7.2% of the 369,000 flowering plant species (the number is disputed) known to science have been assessed on the IUCN Red List. (Bachman *et al*. 2019; Nic Lughadha *et al*. 2016; 2017). Thanks to the Global Tree Assessment (BGCI 2021) many of the world’s tree species have now been assessed. The State of the World’s Trees concluded that the highest proportion of threatened tree species is found in Tropical Africa, and that Cameroon has the highest number (414) of threatened tree species of all tropical African countries (BGCI 2021). This will be further increased by the addition of *Monanthotaxis bali* and will require the updating of the Red Data Book of Cameroon Plants (Onana & Cheek 2011). However, the vast majority of plant species still lack assessments on the Red List (Nic Lughadha *et al*. 2020).

It is imperative to uncover the existence of previously unknown species as soon as possible and to formally name them. Until this is done, they are invisible to science and the possibility of them being assessed for their conservation status and appearing on the IUCN Red List is greatly reduced (Cheek *et al*. 2020b), limiting the likelihood both that they will be proposed for conservation measures, and that such measures will be accepted.

Bali Ngemba, while a Forest Reserve, is not formally protected for nature conservation and is its natural forest habitat is being encroached for agriculture. Protection of Bali Ngemba, with the support of local communities supported by designation as a TIPA (Darbyshire *et al*. 2017) is essential if the 12 endemic and near-endemic plant species unique (or nearly unique) such as *Monanthotaxis bali* are not to become extinct globally, and if the 34 threatened species of that forest are not to move closer to extinction.

## Acknowledgements

This paper was completed as part of the Cameroon TIPAs (Tropical Important Plant Areas) project at RBG, Kew, which is supported by Players of Peoples Postcode Lottery. We thank Lydia Burns and Penny Applebe of Kew’s Foundation for making this possible. The specimen cited in this paper was collected by or with the support of volunteers and sponsored scientists arranged by Earthwatch Europe, Oxford, and we were also assisted greatly by our colleagues such as the late Martin Etuge, but also Innocent Wultoff and Terence Atem. John DeMarco of the then Bamenda Highlands Forest Project persuaded us to make a botanical survey of Bali Ngemba forest in the Bamenda Highlands, and with Anne Gardner supported our visits. The late Dr Benoît Satabié, Drs Gaston Achoundong, Florence Ngo Ngwe, Eric Nana, Jean Lagarde Betti, the current and former directors, of IRAD-National Herbarium of Cameroon, Yaoundé, and their staff are thanked for expediting the collaboration between our two institutes. Janis Shillito typed the manuscript. Two anonymous reviewers are thanked for constructively reviewing an earlier version of this paper.

The authors declare that they have no conflict of interest.

